# Comparative genomics and phylogenomics of *Campylobacter* unveil potential novel species and provide insights into niche segregation

**DOI:** 10.1101/2022.10.11.511782

**Authors:** Sarah Henaut-Jacobs, Hemanoel Passarelli-Araujo, Thiago M. Venancio

## Abstract

*Campylobacter* is a bacterial genus associated with community outbreaks and gastrointestinal symptoms. Studies on *Campylobacter* generally focus on specific pathogenic species such as *C. coli* and *C. jejuni*. Currently, there are thousands of publicly available *Campylobacter* genomes, allowing a more complete assessment of the genus diversity. In this work, we report a network-based analysis of all available *Campylobacter* genomes to explore the genus structure and diversity, revealing potentially new species and elucidating genus features. We also hypothesize that the previously established Clade III of *C. coli* is in fact a novel species (referred here as *Campylobacter spp12*). Finally, we found a negative correlation between pangenome fluidity and saturation coefficient, with potential implications to the lifestyles of distinct *Campylobacter* species. Since pangenome analysis depends on the number of available genomes, this correlation could help estimate pangenome metrics of *Campylobacter* species with less sequenced genomes, helping understand their lifestyle and niche adaptation. Together, our results indicate that the *Campylobacter* genus should be re-evaluated, with particular attention to the interplay between genome structure and niche segregation.

## INTRODUCTION

*Campylobacter* is a Gram-negative Epsilonproteobacteria genus of motile bacteria that can infect humans and other animals. *Campylobacter* species are the leading cause of bacterial foodborne infections, especially transmitted via the consumption of undercooked chicken meat, contaminated water, or dairy products^1^. The genus comprises 38 validly published and non-synonymous species and 7 subspecies according to LPSN (List of Prokaryotic names with Standing in Nomenclature)^2^ (on March 2023).

Among all known *Campylobacter* species, *C. coli* and *C. jejuni* are responsible for over 90% of human campylobacteriosis^1^. The ease of adaptation of these two species to multiple hosts led the World Health Organization to define *Campylobacter* as a pathogen of critical priority for the study and development of new antibiotic treatments^3^. Because of their clinical relevance, *C. coli* and *C. jejuni* have the largest number of publicly available genomes among *Campylobacter* species, creating an unintended bias for genus-wide population structure and ecological studies.

*Campylobacter* species occupy a wide range of niches^4^. For example, *C. coli* is differentiated into three clades (I, II, and III)^5^. While isolates from Clade I are usually associated with acute diarrhea in humans, those belonging to clades II and III are mainly found in environmental sources (e.g., lakes)^5,6^. In contrast, species such as *C. jejuni* present a more clonal population structure with host-specific clonal complexes (e.g., CC-42 and CC-61), which are associated with cattle and sheep^7^. Importantly, these niche preferences can be related to genome sequences, with bacteria with similar lifestyles sharing a greater number of genes. While 16S rRNA sequencing has been widely used to study microbial communities, it is often unable to discriminate closely related genomes and can exhibit bias towards certain species^8^. The development of second and third generation sequencing technologies fueled the use of whole-genome based methods, such as Average Nucleotide Identity (ANI) and genome distance estimation^9^. Here, we employ a network-based approach using genome-wide comparative metrics to explore *Campylobacter* diversity and taxonomic classification.

## Results

### Dataset collection and type strains

We retrieved 3,319 *Campylobacter* genomes from the RefSeq database^10^ in August 2020. After removing genomes with completeness below 90% and duplication levels (a proxy for contamination) greater than 5%, we created a dataset with 3261 *Campylobacter* genomes (Table S1). According to the NCBI classification, 46.24% and 29.9% of the genomes belong to *C. jejuni* and *C. coli*, respectively (Table S1), highlighting an expected overrepresentation of these species. These results also uncovered possible misclassification problems, since overrepresented species might overshadow less studied ones. The LPSN comprises 45 validly named and published *Campylobacter* representative genomes (Table S2), encompassing 38 species and 7 subspecies. We conducted an ANI analysis with all type strain genomes and confirmed that all pairs of type strains from different species had ANI values lower than 95% (the threshold to define bacterial species^11^). We also identified three subspecies with ANI lower than 95% with their respective species’ type strains (*C. fetus* subsp. *testudinum, C. lari* subsp. *Concheus* and *C. pinnipediorum* subsp. *Caledonicus*), suggesting that the communities containing these subspecies might be from distinct novel species. These subspecies were further evaluated using a network-based approach, as described below.

### Network analysis identifies potentially new *Campylobacter* species

We conducted a network analysis based on Mash distances (see methods for details). In this network, nodes and edges represent genomes and the estimated identities (1 – Mash; minimum threshold of 95%), respectively. Each community in the resulting network is considered a different species, as described elsewhere^9^. Species networks inferred with genomic distance or identity are highly structured, with few overlapping communities^9,12^. In our *Campylobacter* network, we found 65 genome communities (Figure 1), which is above the expected number considering the 34 type genomes listed in LPSN. Out of the 65 communities, only 36 contained type strain genomes. We found no communities with more than one species type genome, while two subspecies formed separated communities (*C. lari* subsp. *concheus* and *C. fetus* subsp. *testudinum*). The communities with a type genome (from species or subspecies) were named after their, propagating this classification to all the genomes of the community.

**Figure 1.**
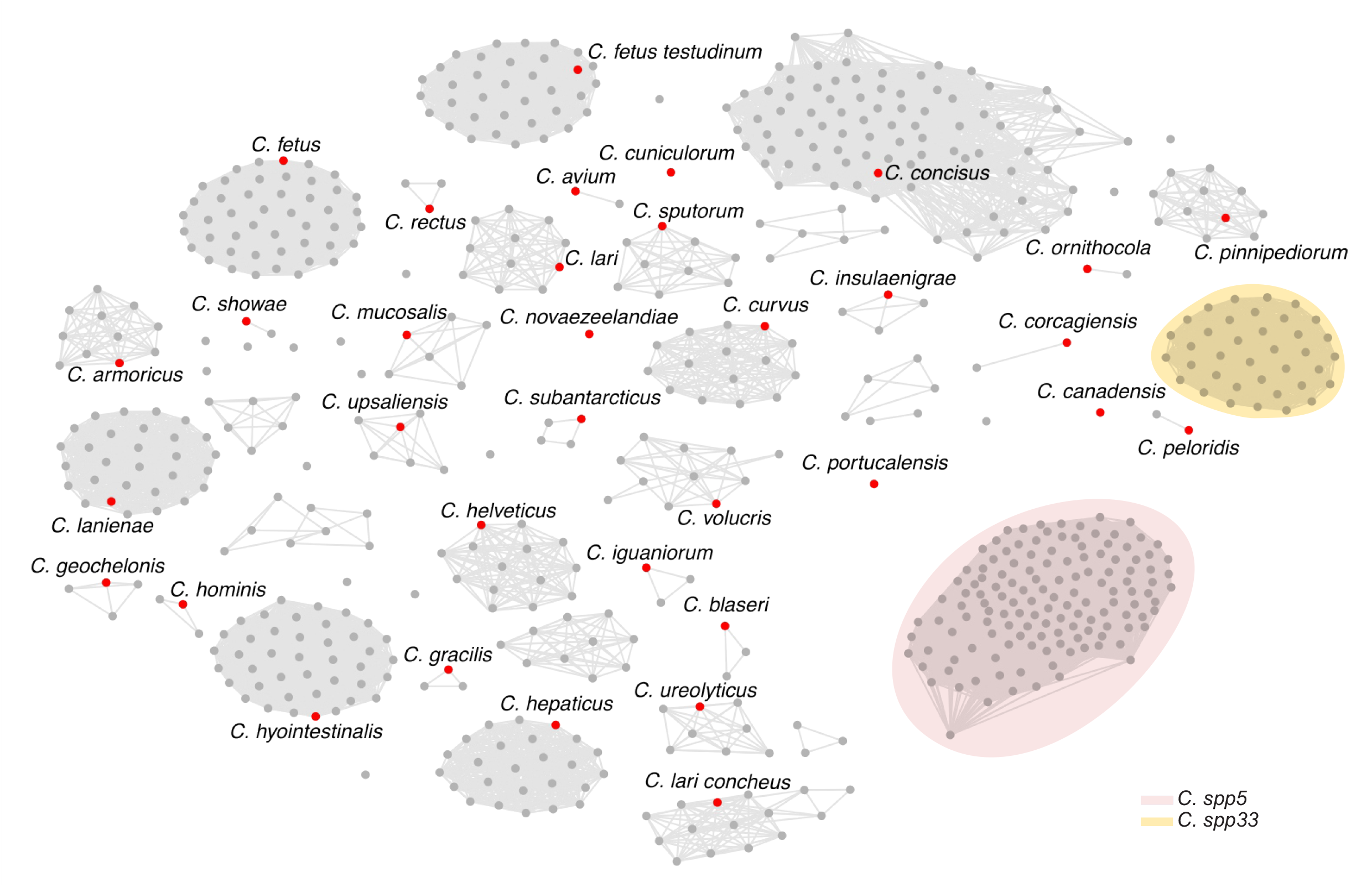
*Campylobacter* network structure. Nodes represent genomes and edges connect those with at least 95% identity. Type strains are highlighted in red. Communities without type strains represent potentially novel species. *C. jejuni* and *C. coli* communities have a large number of genomes, which can make it more difficult to understand the network as a whole, they were omitted to improve clarity.

We named the 29 communities without a type genome as *Campylobacter “sppX”*, where X represents an arbitrary community number (see methods). We found three large unnamed communities: *Campylobacter spp5* (134 genomes), corresponding to the *C. concisus* GSII; *Campylobacter spp12* (78 genomes), representing the well-known *C. coli Clade III*; and *Campylobacter spp33* (37 genomes), comprising genomes from isolates exclusively obtained from swine feces^13^ that form a sister group with *C. lanienae* (Figure 2). These results support that a significant part of the *Campylobacter* diversity remains largely unexplored.

**Figure 2.**
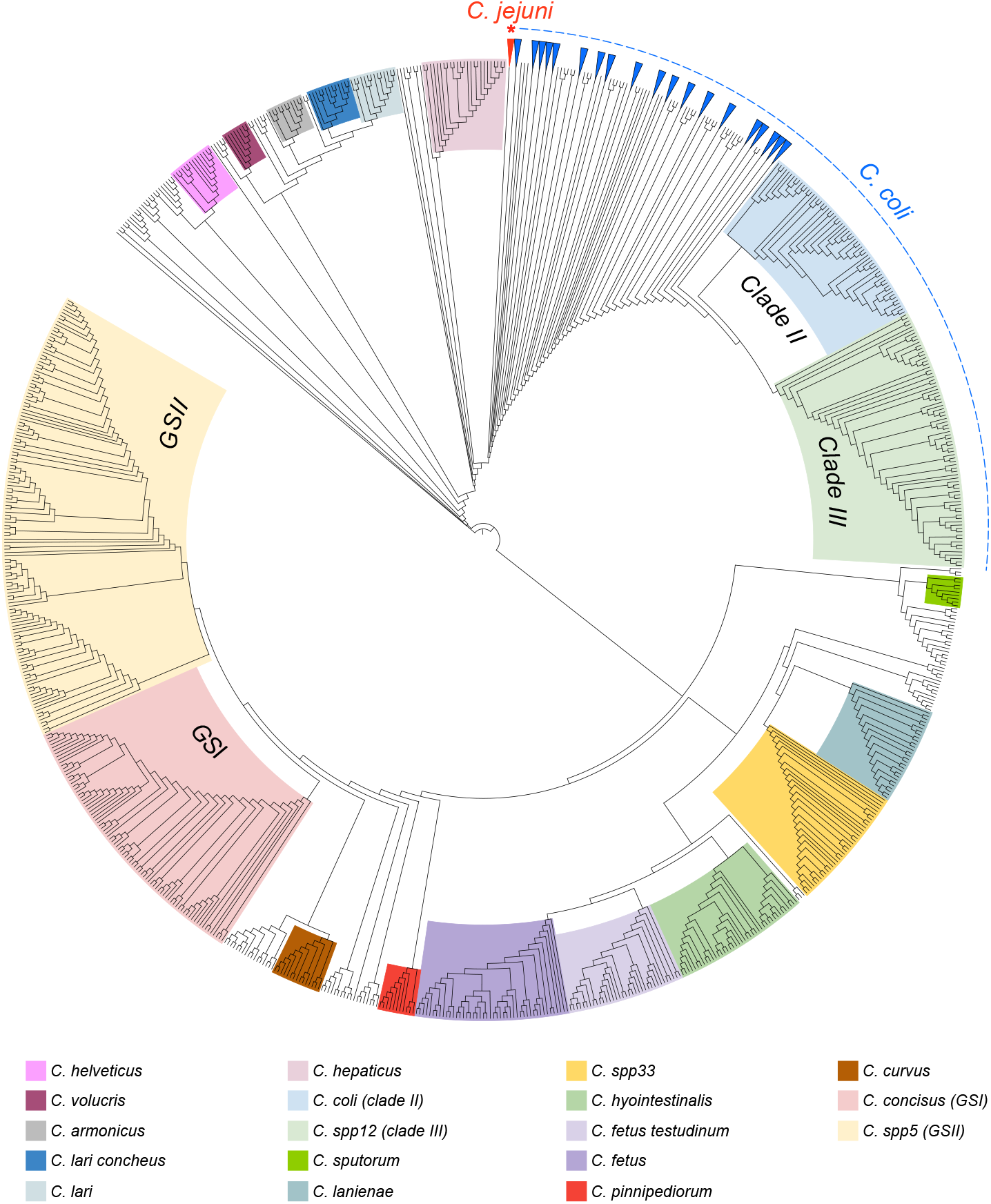
Mash distance tree of *Campylobacter*. Main *Campylobacter* groups are represented in different colors. Branches with many taxa were collapsed to improve visibility.

Besides investigating potentially new species, we also constructed a distance tree to infer the proximity of their respective communities to type species (Figure 2). We observed that *C. coli* is the sister group of *C. jejuni*, as previously described^5^. The *C. coli* population is structured in three main clades^14^. We observed that the *C. coli* Clade III is an independent community in the distance network and is paraphyletic in relation to the other *C. coli* genomes (Figure 2). Although *C. coli* Clade II is a monophyletic sister group of Clade I in the distance tree, the high number of network connections with other *C. coli* genomes does not support its separation as an independent community.

We also corroborated the split of *C. concisus* into two distinct genomospecies (GSI and GSII), as previously proposed^15^. Within the 216 *C. concisus* genomes available in RefSeq, 82 belong to *C. concisus* GSI, while 134 genomes formed a distinct community named as *Campylobacter spp5* (*C. concisus* GSII). The two communities also differ in GC content (37.22% in *C. concisus* and 39.19% *in Campylobacter spp5*; Wilcoxon test, p-value < 0.001) and genome sizes (1.9 Mbp in *C. concisus* and 2.0 Mbp in *Campylobacter spp5*; Wilcoxon test, p-value < 0.001), supporting their split into two different species. Accordingly, it has been proposed that the GC content within *Campylobacter* species should not vary by more than 1%^16^.

The distance tree also provides insights into novel species and niche segregation. For example, *C. fetus* subsp. *testudinum* is probably a new species, typically found in mammals. Furthermore, we also observed a clear division of *C. lari* into two clades, corresponding to the two subspecies previously proposed based on 16S rRNA: *C. lari* subsp. *Lari* and *C. lari* subsp. *concheus*^17^. These results imply that the *C. lari* phylogeny also warrants revisions.

### *Campylobacter* phylogeny of representative genomes and species reclassification

We used all core single-copy genes from 65 representative genomes (Table 2) to reconstruct the *Campylobacter* phylogenetic tree. This phylogenetic reconstruction uncovered a remarkable difference in GC content across clades. (Figure 3). For example, the clade comprising *C. showae* and *C. rectus* have GC content greater than 40%, far greater than the 30.4% found in the *C. jejuni* clade.

**Figure 3.**
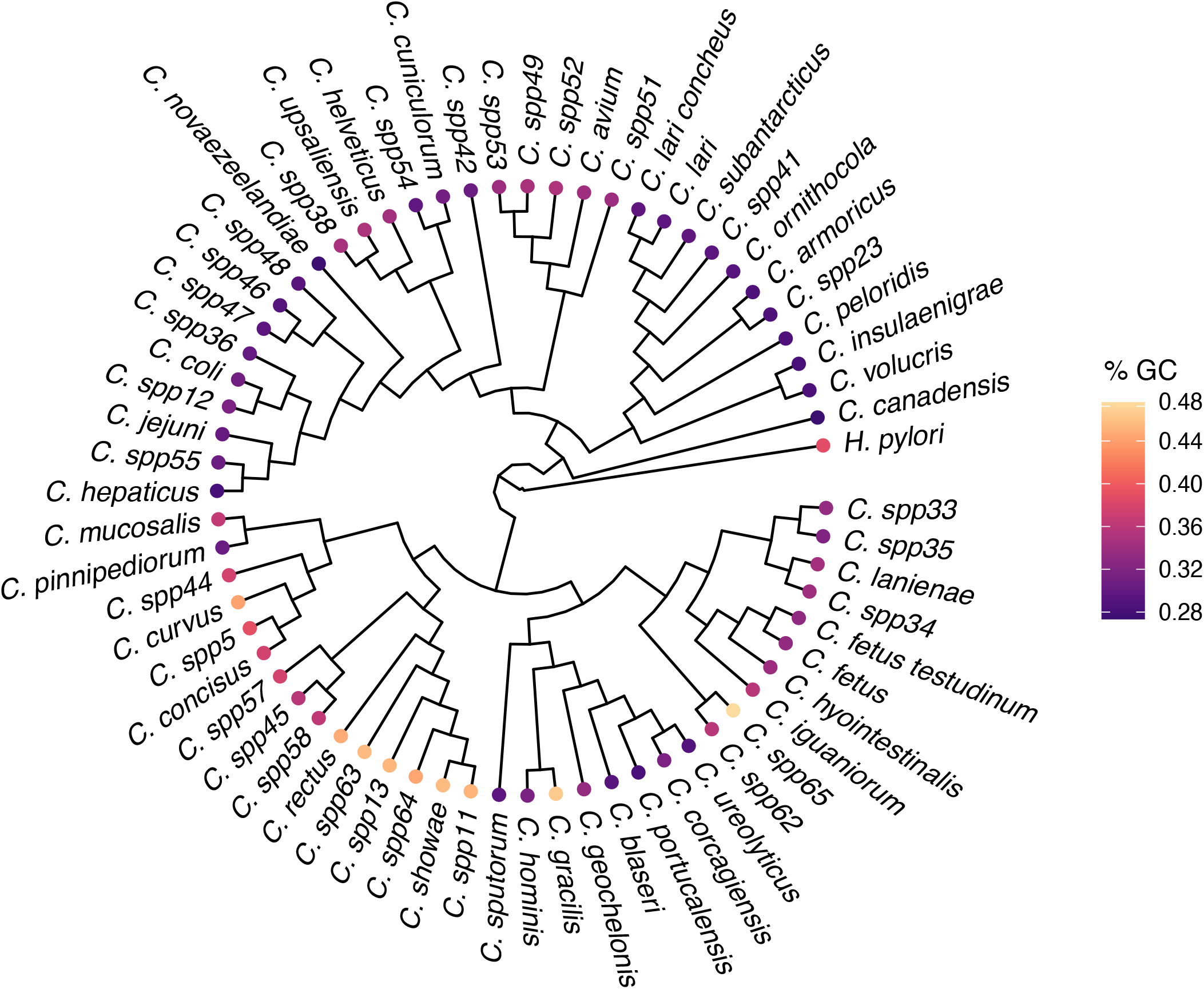
Phylogenetic reconstruction and GC content in *Campylobacter*. The Maximum likelihood tree was reconstructed using core single-copy genes from one representative genome from each of the 65 *Campylobacter* communities. *H. pylori* was used as outgroup.

We also used the representative genomes to estimate a misclassification rate of 7.57% in *Campylobacter spp* (247 misclassified genomes; Table S3). We considered as misclassified only those genomes previously annotated at the species level and re-assigned either to a different validly published species or to a new community (*Campylobacter sppX*). Therefore, genomes identified as “*Campylobacter sp*.” were not considered misclassified if assigned to a community.

We found extreme misclassification levels in eight known species (Table 1), particularly *C. showae* (81.82%, n=11). Given the small number of *C. showae* genomes, it would be premature to reach a conclusion on this species. However, as most of the genomes formed a cluster apart from the type species, we hypothesize that the type genome may not satisfactorily represent the group. Species such as *C. concisus* and *C. coli* showed significant reclassification levels due to their known heterogeneity^5,15^ (61.86%, n=215 and 7.96%, n=980, respectively). Further, genomes submitted to the RefSeq database as *C. aviculae, C. estrildidarum, C. taeniopygiae*, and *C. troglodytis* were marked as reclassified because of the absence of their respective representative genomes in LPSN at the time of the analysis. Finally, 92 genomes previously identified as *Campylobacter sp*. (54.76%). were assigned to communities composed by known species, particularly *C. jejuni*.

**Table 1:** Rate of misclassified genomes per species in the RefSeq database. Genomes originally assigned as belonging to a species and appeared an unexpected community (different from that of their respective type strain genome) were considered misclassified.

| Original classification | Reclassification rate (%) |
| --- | --- |
| *C. coli* | 7.96 (78/980) |
| *C. concisus* | 61.86 (133/215) |
| *C. fetus* | 27.27 (10/35) |
| *C. hepaticus* | 3.85 (1/26) |
| *C. jejuni* | 0.39 (6/1551) |
| *C. lanienae* | 6.45 (2/31) |
| *C. lari* | 50.00 (9/18) |
| *C. showae* | 81.82 (9/11) |

**Table 2:** Representative genomes of each community detected in this study. The communities were named after their type strains. Communities without type strains were arbitrarily named with a sequential suffix and received a random representative genome.

| Community name | Refseq accession | Community name | Refseq accession |
| --- | --- | --- | --- |
| *C. armoricus* | GCF_013372105.1 | *Campylobacter spp57* | GCF_012978795.1 |
| *C. avium* | GCF_002238335.1 | *Campylobacter spp58* | GCF_012978835.1 |
| *C. blaseri* | GCF_013201895.1 | *Campylobacter spp62* | GCF_013416015.1 |
| *C. canadensis* | GCF_013177655.1 | *Campylobacter spp63* | GCF_900573945.1 |
| *C. coli* | GCF_003590975.1 | *Campylobacter spp64* | GCF_900573985.1 |
| *C. concisus* | GCF_001298465.1 | *Campylobacter spp65* | GCF_902795495.1 |
| *C. corcagiensis* | GCF_013201645.1 | *C. sputorum* | GCF_008802035.1 |
| *C. cuniculorum* | GCF_002104335.1 | *C. subantarcticus* | GCF_000816305.1 |
| *C. curvus* | GCF_013372125.1 | *C. upsaliensis* | GCF_900637395.1 |
| *C. fetus* | GCF_900475935.1 | *C. ureolyticus* | GCF_013372225.1 |
| *C. fetus testudinum* | GCF_002973655.1 | *C. volucris* | GCF_008801985.1 |
| *C. geochelonis* | GCF_013201685.1 |  |  |
| *C. gracilis* | GCF_001190745.1 |  |  |
| *C. helveticus* | GCF_002080395.1 |  |  |
| *C. hepaticus* | GCF_001687475.2 |  |  |
| *C. hominis* | GCF_900446395.1 |  |  |
| *C. hyointestinalis* | GCF_013372145.1 |  |  |
| *C. iguaniorum* | GCF_000736415.1 |  |  |
| *C. insulaenigrae* | GCF_900637885.1 |  |  |
| *C. jejuni* | GCF_000254515.1 |  |  |
| *C. lanienae* | GCF_002139935.1 |  |  |
| *C. lari* | GCF_013372185.1 |  |  |
| *C. lari concheus* | GCF_008245025.1 |  |  |
| *C. mucosalis* | GCF_013372205.1 |  |  |
| *C. novaezeelandiae* | GCF_004323845.1 |  |  |
| *C. ornithocola* | GCF_013201605.1 |  |  |
| *C. peloridis* | GCF_000816785.1 |  |  |
| *C. pinnipediorum* | GCF_002021925.1 |  |  |
| *C. portucalensis* | GCF_009690845.1 |  |  |
| *C. rectus* | GCF_004803795.1 |  |  |
| *C. showae* | GCF_004803815.1 |  |  |
| *Campylobacter spp11* | GCF_000313615.1 |  |  |
| *Campylobacter spp12* | GCF_001489835.1 |  |  |
| *Campylobacter spp13* | GCF_000344295.2 |  |  |
| *Campylobacter spp23* | GCF_000816245.1 |  |  |
| *Campylobacter spp33* | GCF_900446565.1 |  |  |
| *Campylobacter spp34* | GCF_002139105.1 |  |  |
| *Campylobacter spp35* | GCF_002139895.1 |  |  |
| *Campylobacter spp36* | GCF_002179165.1 |  |  |
| *Campylobacter spp38* | GCF_002738235.1 |  |  |
| *Campylobacter spp41* | GCF_004357905.1 |  |  |
| *Campylobacter spp42* | GCF_004378855.1 |  |  |
| *Campylobacter spp44* | GCF_012978635.1 |  |  |
| *Campylobacter spp45* | GCF_004803855.1 |  |  |
| *Campylobacter spp46* | GCF_005406205.1 |  |  |
| *Campylobacter spp47* | GCF_005406215.1 |  |  |
| *Campylobacter spp48* | GCF_005406225.1 |  |  |
| *Campylobacter spp49* | GCF_010092865.1 |  |  |
| *Campylobacter spp5* | GCF_902460635.1 |  |  |
| *Campylobacter spp51* | GCF_006864365.1 |  |  |
| *Campylobacter spp52* | GCF_006864425.1 |  |  |
| *Campylobacter spp53* | GCF_006864455.1 |  |  |
| *Campylobacter spp54* | GCF_008633875.1 |  |  |
| *Campylobacter spp55* | GCF_009362725.1 |  |  |

We also compared the results obtained by the genome analysis with the *Campylobacter* classification provided by GTDB (Release 207, downloaded in August 2022). GTDB has a standardized framework to classify genomes in higher taxonomic ranks (e.g., genus)^18^ by following a phylogenomic approach based on a set of conserved single-copy proteins. This scheme divides *Campylobacter* in five different genera (Table S1): *Campylobacter, Campylobacter A, Campylobacter B, Campylobacter D*, and *Campylobacter E*. Importantly, this distribution is coherent with the phylogenetic reconstruction and community analysis described here. The largest genus, *Campylobacter E*, comprised the *C. coli* and *C. jejuni* communities, whereas the second largest genus, *Campylobacter A*, harbored the monophyletic group of *C. concisus, Campylobacter spp5, C. curvus*, and *C. pinnipediorum*.

A comparison with the TYGS (Type Strain Genome Server)^19,20^ was also conducted using the representative genomes of each community (Table S4). TYGS corroborated our findings, with most of the unknown communities being identified as potentially new species. We also found that *Campylobacter spp55* (n=1), and *Campylobacter spp38* (n=2) were identified as *C. bilis* and *C. vulpis* respectively, both found in the most recent LPSN update^2^ (Table S2). Finally, the TYGS classification of representative genomes of the most well studied communities were consistent with our observations.

### Pangenome analysis

The pangenome analysis of a bacterial species provides insights into its lifestyle^21^. A pangenome is the total set of non-redundant genes in a species. It comprises the core genome, the set genes present in all isolates; the accessory genome, with genes present in more than one but not in all isolates; and the isolate-exclusive genes^22^. We computed the genomic fluidity and pangenome openness to investigate the overall architecture of the *Campylobacter* pangenome (Table 3, Figure 4). The genomic fluidity measures how dissimilar genomes are at the gene level^23^, while pangenome openness reflects the propensity to detect new genes, as new genomes are included in the analysis. Since sample size can bias pangenome estimation, we analyzed only those communities with more than 20 genomes.

**Table 3:**
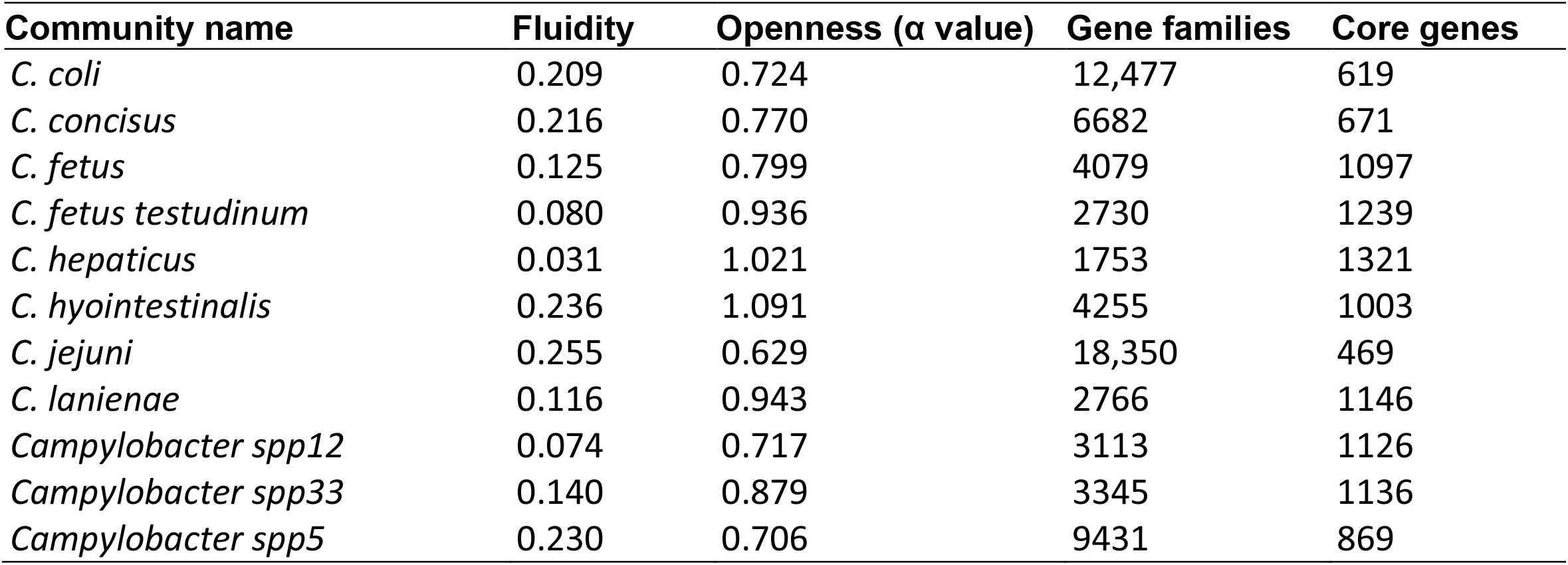
Pangenome features for each community with 20 or more genomes.

**Figure 4.**
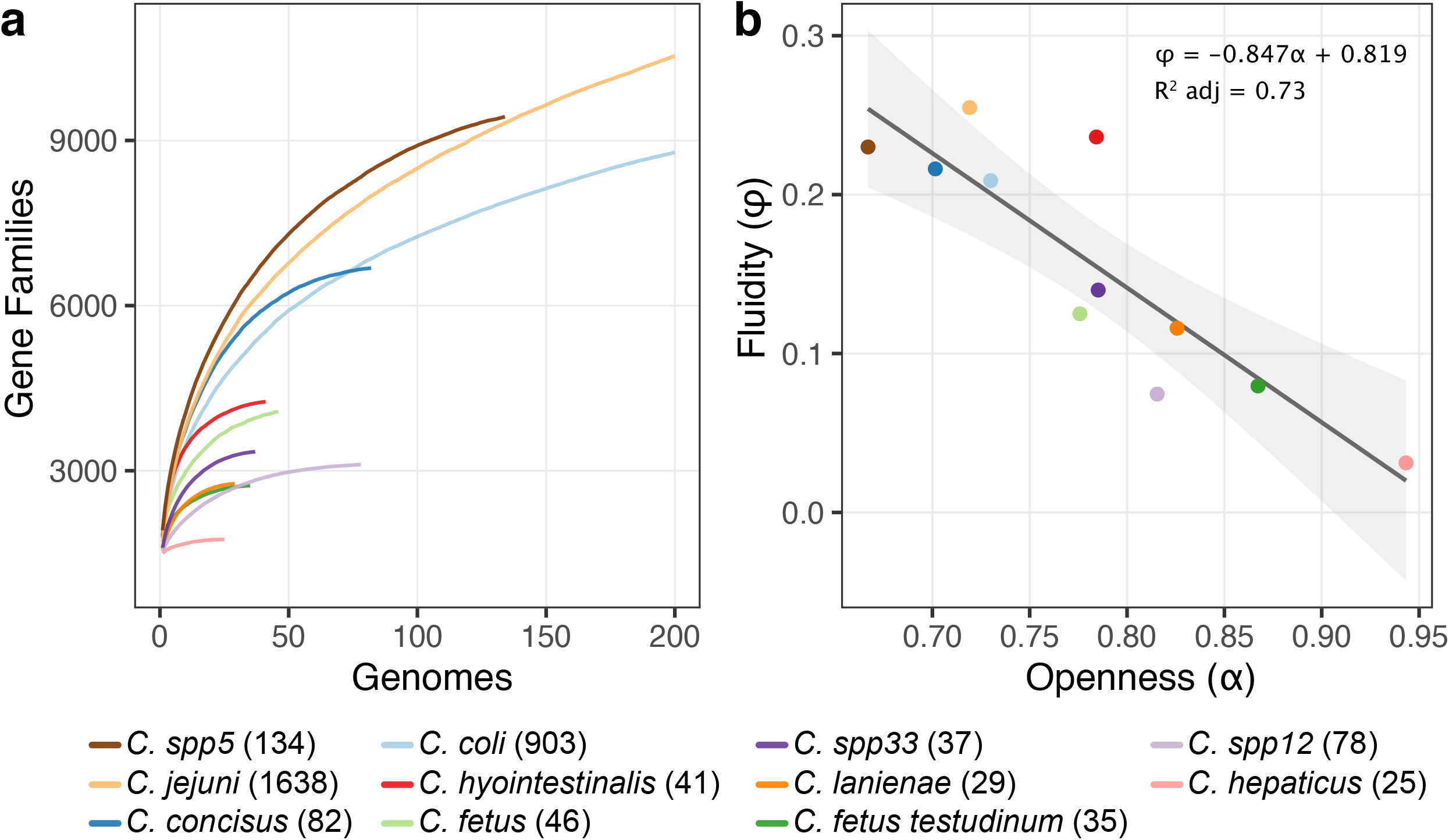
Pangenome openness and fluidity of *Campylobacter* species. **a)** Pangenome cumulative curve from eleven *Campylobacter* species from communities with at least 20 genomes. The number of genomes of each species are indicated within parentheses. Due to the high number *C. jejuni* and *C. coli* genomes, we limited the x-axis to 200 genomes. **b)** Simple linear regression to predict *Campylobacter* pangenome fluidity (ϕ) based on pangenome openness (α). This analysis was conducted with the same communities from panel a.

Previous analyses on the connection between pangenome openness and genomic fluidity have shown that these two metrics are not necessarily correlated, with both pangenome fluidity and saturation being related to its openness^24^. We found that genomic fluidity of *Campylobacter* is negatively correlated with saturation coefficient (α) (*ρ* = -0.8733; p-value = 0.0004), with greater values of α representing a closer pangenome. We found a significant regression equation (F_1;9_ = 28.927; p-value < 0.0001), with an *R*^*2*^ of 0.73 (Figure 4b) and predicted genomic fluidity of 0.8187 − 0.8467*α*, meaning that the genomic fluidity decreased 0.8467 units for each unit of the parameter α. This result provides a model for estimating the genomic fluidity of species without enough genomes to be included in the pangenome analysis.

We observed that *C. hepaticus* is genetically conserved and homogeneous. This species has the lowest diversity and the highest saturation coefficient (closed pangenome) (Figure 4a) across *Campylobacter* communities, supporting a low genomic diversity and reduced HGT in this species. However, it is important to mention that many *C. hepaticus* genomes are from the same bioproject, which might result in unrealistic diversity estimates for this species. On the other hand, species such as *C. jejuni* and *C. coli* have a high genomic fluidity, which is in line with the generalist ecological character of both species, with isolates from various host sources.

## DISCUSSION

The *Campylobacter* genus is often associated with gastrointestinal diseases and antibiotic resistance, having a strong study bias towards clinically relevant species such as *C. jejuni* and *C. coli*. Here, we analyzed 3,261 *Campylobacter* genomes from several species and sources. We identified 65 different communities within *Campylobacter*, way over the 38 species originally listed on the LPSN. We found a portion of these novel communities (here considered as potential new species) to be closely related to some *Campylobacter* groups such as the *C. lari* and *C. fetus* groups. Our ANI-based network analysis also showed that 7.57% of the *Campylobacter* genomes warrants phylogenetic reclassification.

*C. concisus* is an oral bacterium that consists of two genomospecies (GSI and GSII) that are undistinguishable at the phenotypic level^15^. Our results support this division, as we identified two *C. concisus* communities, the first containing the species’ type strain with 82 genomes, and *Campylobacter spp5* with 134 genomes. Similarly, we also found support to separate *C. coli* in two communities: one comprising the original type strain and other 902 genomes, and a second community with 78 genomes (referred as *Campylobacter spp12*).

We also reported strong evidence for potentially novel species into the *C. fetus* group. *Campylobacter spp33* comprises 37 genomes of isolates obtained only from swine feces^13^, supporting a specific niche occupation. Interestingly, specific niche occupation is a common feature of the *C. fetus* group ^25^. We can further divide this clade using characteristics like the low capacity to cause diseases in humans (*C. lanienae-*related species) and selenium metabolism deficiency (*Campylobacter spp33* clade^13^). Finally, the identification of *Campylobacter spp33* as a novel species should be further investigated for a more thorough and formal description.

We found that genomic fluidity and pangenome saturation coefficient are negatively correlated in the *Campylobacter* genus. Although these two parameters are usually conceptually associated, previous findings have shown that there was not a direct and unequivocal correlation between their values, but that fluidity was just one of the determinants of genome openness^24^. *C. hepaticus* showed a close and homogeneous pangenome, although one must consider that a high fraction of the *C. hepaticus* genomes were from the same study, which could result in the underestimation of the *C. hepaticus* diversity. Finally, it is worth mentioning that more in-depth investigations of the large unnamed communities described here should be conducted to validate them as novel species.

## METHODS

### Dataset collection

We downloaded 3,319 *Campylobacter* genomes available in RefSeq using Datasets v4.2.0 (https://www.ncbi.nlm.nih.gov/datasets/) in August 2020. We used BUSCO v4.1.0^26^ and removed the genomes with completeness lower than 90%, and duplication greater than 5%. Genomic metadata were gathered from RefSeq using the Reutils package v. 0.2.3 (http://mirrors.ucr.ac.cr/CRAN/web/packages/R.utils/R.utils.pdf).

All the genomes were analyzed using Mash v1.1^27^. We compared different Mash sketch values to find the parameter with the lowest error rate^9^. We used the mean distances generated by sketch sizes of 1000, 2000, and 5000 bp, removing genome distances greater than 0.8. We used the reciprocal of Mash distance to estimate the identity (1 – Mash). For type strain analyses, we computed ANI using pyani v0.2.10^28^.

The network analyses of the distances between the genomes were conducted using igraph v1.2.7^29^. We generated a weighted graph with nodes and edges representing genomes and genetic identities between them, respectively. Only edges representing identities greater than 95% were retained.

### Type strains validation

The pre-set *Campylobacter* type strains were retrieved from LPSN^2^ in March 2023. The valid accession numbers of each species and subspecies were manually gathered, and we used the information in LPSN to identify validly reported and documented type genomes (38 representing valid species, 7 representing valid subspecies) (Table S2). Next, we conducted an ANI analysis with Pyani v0.2.10^28^ to verify that all the genome pairs are at least 95% identical to each other.

### Network analysis

We employed the label propagation algorithm to detect communities in the genome distance network, as previously reported^9^. Communities with type genomes were named after them, while communities without a type genome were arbitrarily identified as *Campylobacter sppX*, where X represents the number of the community. These communities had one of its members randomly assigned as its representative genome. The named communities were contrasted with the RefSeq taxonomic classification to estimate the level of misclassification.

### Phylogeny and pangenome characterization

After selecting a representative genome from each *Campylobacter* community, we used Orthofinder v.2.5.2^30^ to infer orthologs. Single-copy genes were used to reconstruct the phylogenetic tree using IQ-TREE v.1.6.12^31^ with the TEST argument to find the best fitting model. One thousand bootstrap replicates were generated to assess the significance of internal nodes. The *Helicobacter pylori* strain ATCC 43504 (Accession number: GCF_900478295.1) were used as outgroup. We used ggtree^32^ to visualize the reconstructed tree. GTDB data (July 2022) and TYGS data (March 2023) were used to analyze different phylogenetic groups within *Campylobacter*^19,20,33^.

We conducted the pangenome analysis for those communities with more than 20 genomes with Roary v3.13.0^34^. Micropan v2.1^35^ and Pagoo v0.3.9^36^ were used to assess pangenome fluidity values and openness values, respectively.

## Author contributions

Conceived the study: HP-A and TMV. Data analysis: HP-A, SH-J. Funding, project coordination and infrastructure: TMV. Manuscript writing: HP-A, SH-J, TMV.

## Acknowledgments

This work was supported by Fundação Carlos Chagas Filho de Amparo à Pesquisa do Estado do Rio de Janeiro (FAPERJ; grants E-26/203.309/2016 and E-26/203.014/2018), Coordenação de Aperfeiçoamento de Pessoal de Nível Superior-Brasil (CAPES; Finance Code 001), and Conselho Nacional de Desenvolvimento Científico e Tecnológico (CNPq). The funding agencies had no role in the design of the study and collection, analysis, and interpretation of data and in writing.

